# A rhamnose-inducible system for precise and temporal control of gene expression in cyanobacteria

**DOI:** 10.1101/227355

**Authors:** Ciarán L. Kelly, Andrew Hitchcock, Antonio Torres-Méndez, John T. Heap

## Abstract

Cyanobacteria are important for fundamental studies of photosynthesis and have great biotechnological potential. In order to better study and fully exploit these organisms, the limited repertoire of genetic tools and parts must be expanded. A small number of inducible promoters have been used in cyanobacteria, allowing dynamic external control of gene expression through the addition of specific inducer molecules. However, the inducible promoters used to date suffer from various drawbacks including toxicity of inducers, leaky expression in the absence of inducer and inducer photolability, the latter being particularly relevant to cyanobacteria which, as photoautotrophs, are grown under light. Here we introduce the rhamnose-inducible *rhaBAD* promoter of *Escherichia coli* into the model freshwater cyanobacterium *Synechocystis* sp. PCC 6803 and demonstrate it has superior properties to previously reported cyanobacterial inducible promoter systems, such as a non-toxic, photostable, non-metabolizable inducer, a linear response to inducer concentration and crucially no basal transcription in the absence of inducer.

## INTRODUCTION

Photoautotrophic microorganisms have great potential for the sustainable production of chemicals from carbon dioxide using energy absorbed from light. Cyanobacteria including *Synechocystis* sp. PCC 6803 (‘*Synechocystis*’ hereafter) and *Synechococcus* sp. PCC 7002 have been successfully engineered to produce 2,3-butanediol^1,2^, lactate^3^, isobutanol^4^, plant terpenoids^5^ and ethanol^6–9^, and to allow the utilisation of xylose^10^. Cyanobacteria, particularly *Synechocystis,* are also used as model organisms for fundamental studies of important processes such as photosynthesis^11–16^, circadian rhythms^17–19^ and carbon-concentrating mechanisms^20–23^. Due to specific challenges, genetic modification of cyanobacteria is more difficult than genetic modification of model heterotrophic microorganisms such as *Escherichia coli* and *Saccharomyces cerevisiae*. These challenges include polyploidy^24,25^, which makes the isolation of segregated recombinant strains slow and laborious; genetic instability of heterologous genes^26^ and limited synthetic biology tools and parts such as promoters and expression systems. Improved synthetic biology capabilities for cyanobacteria would be useful for both fundamental and applied studies.

Inducible promoters are important tools which allow flexible control over gene expression, which is useful in many fundamental and applied studies. Unlike the limited number of constitutive promoters which have been shown to function well in cyanobacteria^6,10,27–29^), inducible promoters provide access to a wide, continuous range of gene expression levels using a single genetic construct, simply by varying inducer concentrations^30^. Furthermore, inducible promoters also allow control over the timing of expression of a gene of interest. An ideal inducible promoter system would have certain properties: Firstly, the promoter should not ‘leak’, that is, there should be no basal transcription in the absence of inducer, allowing very low expression levels to be used, and avoiding premature expression during strain construction and segregation, which can be associated with toxicity and mutation^26,31,32^. Secondly, the inducer molecule should be non-toxic, non-metabolisable, readily available and stable under experimental conditions (including under light in the case of photoautotrophic organisms), allowing sustained expression with no impact on growth caused as an artefact of the expression system itself. Thirdly, expression should demonstrate a near-linear response to inducer concentration over a wide range. Finally, expression should have a consistent unimodal distribution across a population of cells.

Several inducible promoter systems have been described in *Synechocystis* spp. and *Synechococcus* spp., but none are ideal. Metal ion-inducible promoters have been described which respond to nickel, copper, cadmium, arsenic and zinc^33–36^. Unfortunately these systems have disadvantages including the presence of many of the metals in standard growth media^37^, a narrow range of useful concentrations because the concentrations required for detectable and unimodal induction are close to toxic levels^34^, and some are ‘leaky’ in the absence of inducer. The use of metals as inducers also has the potential to disrupt metal homeostasis, resulting in the sequestration of metals required as essential cofactors of many enzymes involved in photosynthesis and related metabolic pathways^11,38–40^. Synthetic inducible promoters have also been constructed and used in cyanobacteria. Two promoter systems using the tetracycline-responsive repressor TetR and its cognate operator sites have been engineered for use in cyanobacteria. The first example for use in *Synechocystis* resulted in a well-characterised, anhydrotetracyline (aTc)-responsive promoter with low leakiness and a good dynamic range^41^. Unfortunately the inducer aTc is extremely sensitive to light and therefore induction from this promoter was transient and required high concentrations of aTc. The second example, in *Synechococcus* sp. PCC 7002, suffered the same issues with photolability of the inducer and low expression by comparison to a commonly-used strong constitutive promoter^42^. It is clear therefore that aTc-based inducible promoters are unsuitable for photoautotrophic growth conditions. The non-metabolisable analogue of lactose, isopropyl β-D-1-thiogalactopyranoside (IPTG) has also been tested for use as an external inducer of lac-based promoters in a variety of cyanobacterial strains^1,27,43–45^, with mixed performance in terms of dynamic range and leakiness in absence of inducer. Finally, use of a green-light inducible promoter in *Synechocystis* sp. PCC 6803 has been reported^46^, but isolating the specific wavelengths required for induction from natural or white light used for growth is difficult, leading to unwanted induction.

To-date, heterologous promoters associated with the AraC/XylS family of positive transcriptional regulators have not been used in cyanobacteria. One promising candidate is the L-rhamnose-inducible *rhaBAD* promoter system of *E. coli*, which naturally has almost all of the ideal properties described above^47–49^. Recently this system was optimised in *E. coli* by the identification of L-mannose as a non-metabolisable inducer and constitutive expression of the activating transcription factor RhaS in order to make the system independent of the native regulatory cascade^50^.

Here we introduce the *rhaBAD* promoter of *E. coli* into *Synechocystis* sp. PCC 6803, characterise its behaviour, assess inducer stability and investigate the effects of modifying various promoter sequence elements and of varying expression of the transcription factor RhaS. The result is an inducible expression system with several important advantages over expression systems previously characterised in cyanobacteria including precise control of the strength and timing of induction as well as sustained gene expression in the presence of light. This system is likely to be very useful and widely applicable in *Synechocystis* and other cyanobacteria.

## RESULTS AND DISCUSSION

### Analysis of the *E. coli rhaBAD* promoter in a heterologous *Synechocystis* context

In the heterologous context of a *Synechocystis* cell the *rhaBAD* promoter might be expected to perform differently than in the native *E. coli* host. To assess this, we considered the known functional features of the *rhaBAD* promoter and whether the relevant transcription factors in *Synechocystis* were present, and if present, investigated the conservation of their functionally-important amino acid residues.

The *rhaBAD* promoter (Figure 1) contains three types of operator sequences for the binding of three distinct transcription factors: the cAMP (cyclic adenosine monophosphate) receptor protein (CRP); RhaS, which in *E. coli* mediates transcriptional activation of the *rhaBAD* operon in response to L-rhamnose; and RpoD, the primary vegetative sigma 70 factor of *E. coli* RNA polymerase (RNAP).

**Figure 1.**
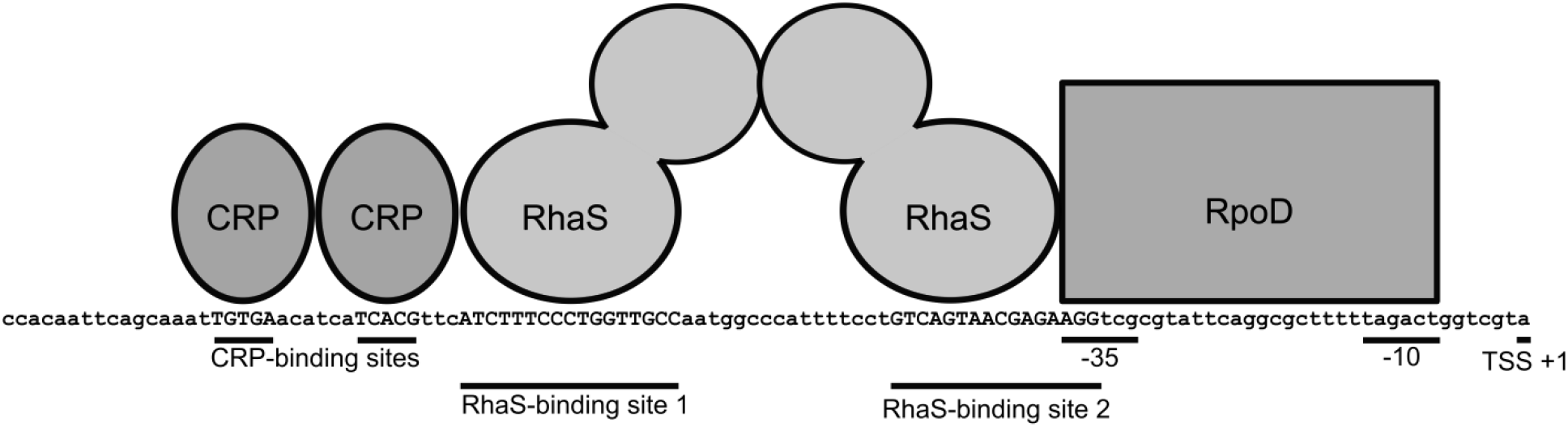
The *rhaBAD* promoter of *E. coli* showing transcription factors and binding sites. Binding site sequences are in uppercase letters and labelled. The −35 and −10 operators^47,51^ are the binding sites for RpoD, the sigma 70 factor of RNA polymerase. TSS +1 is the transcriptional start site.

The genome of *Synechocystis* encodes SYCRP1, a homolog of *E. coli* CRP^52,53^ with 27% identity and 49% similarity to the *E. coli* protein. In *E. coli*, CRP binds to promoters containing specific binding sites when the concentration of cAMP is high, for example when glucose is scarce and other carbon sources must be metabolised for growth. The CRP-binding site in the *rhaBAD* promoter has been shown to be essential for this promoter to function fully in *E. coli*^47,54^. In *Synechocystis*, SYCRP1 has been shown to positively and negatively regulate a number of promoters in response to changing cAMP concentrations^53,55^. The sequence of the CRP-binding site in *Synechocystis* (tgtgaNNNNNNtcaca) differs by only one nucleotide to the CRP-binding site sequence found in the *E. coli rhaBAD* promoter (tgtgaNNNNNNtcac<u>g</u>), which suggests SYCRP1 might bind to this heterologous promoter sequence^56,57^.

To the best of our knowledge, positively-regulated AraC/XylS-type expression systems like those in *E. coli* have not been reported in *Synechocystis* or in other cyanobacteria. In *E. coli*, the positive transcriptional regulator RhaS is essential for transcription from the *rhaBAD* promoter. We used BLASTP^58^ to search the genome of *Synechocystis* for a homolog of *E. coli* RhaS. No protein with significant similarity to *E. coli* RhaS was identified, suggesting that heterologous expression of the *rhaS* gene of *E. coli* which encodes this protein would be required for the heterologous *rhaBAD* promoter from *E. coli* to function in *Synechocystis*.

It has been hypothesised that differences in the RNA polymerase components between cyanobacteria and *E. coli* are one reason for *E. coli* promoters failing to function as expected when used in cyanobacteria^59^. With this in mind, the RpoD sigma factor of *E. coli* and the SigA sigma factor of *Synechocystis* were compared by alignment of their amino acid sequences (Figure 2). RpoD (accession number: NP_417539.1) is the *E. coli* primary vegetative sigma 70 factor, which binds to the −35 and −10 regions of the *rhaBAD* promoter in *E. coli*, and SigA (accession number: ALJ69094.1) is the *Synechocystis* primary sigma factor. The two orthologs share 59% identity and 78% similarity but as the *Synechocystis* protein is much smaller than the *E. coli* ortholog (425 and 613 amino acids respectively), the overall coverage is only 46%, with the N-termini sharing little similarity in contrast to the good alignment at the C-termini, which is the most conserved region across the sigma 70 family of transcription factors^60–62^. The C-termini of sigma 70 factors contain the DNA-binding domains, with conserved and well-defined functional regions^63^. Region 2 is responsible for interaction with the −10 element of the promoter and region 4.2 is responsible for interaction with the −35 element^64,65^. The sequence of the −10 element-binding domain of the *Synechocystis* sigma 70 factor, RTIRLPVH differs only in one amino acid from the *E. coli* sequence RTIRIPVH (Figure 2), which suggests this protein is likely to bind to the −10 element of the *rhaBAD* promoter. Even more encouragingly, the amino acid sequence of the −35 element-binding domain, VTRERIRQIEAKALRKLRHP, is perfectly conserved between both *Synechocystis* and *E. coli* proteins (Figure 2). Finally, it is known that two residues of the *E. coli* RNAP sigma 70 factor protein, RpoD are essential for interaction with two residues of RhaS when both proteins are bound to the DNA^66^. These interactions are formed between R599 of the sigma 70 factor and D241 of RhaS, as well as K593 of the sigma 70 factor and D250 of RhaS. Both of these residues are found in the *Synechocystis* sigma 70 factor protein, corresponding to R412 and K406 respectively (Figure 2). The above analysis suggested that the the *E. coli rhaBAD* promoter is likely to be functional in *Synechocystis*, and will probably require RhaS to be provided.

**Figure 2.**
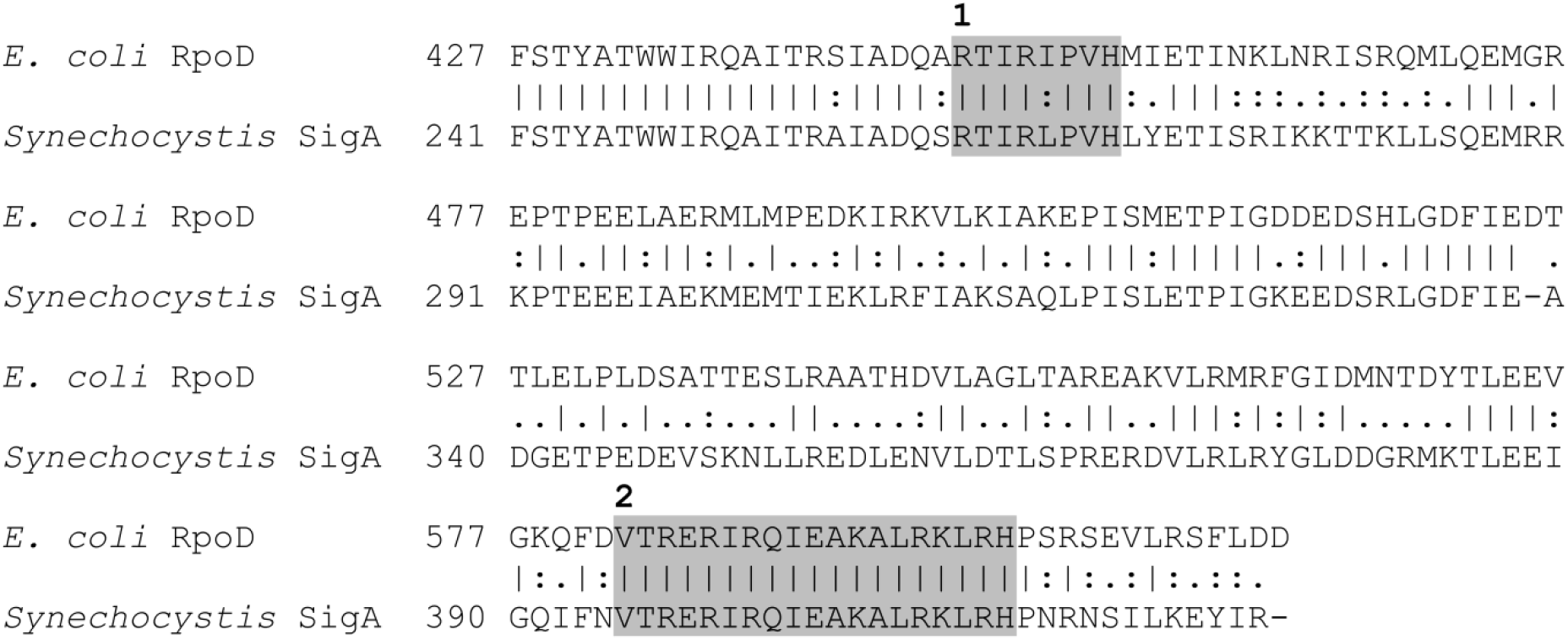
Sequence alignment of RNA polymerase sigma 70 factors from *E. coli* and *Synechocystis*. RpoD (NP_417539.1) of the *E. coli* K12 strain MG1655 was aligned pairwise with SigA (ALJ69094.1) of *Synechocystis* sp. PCC 6803 using EMBOSS Needle^67^ accessed at https://www.ebi.ac.uk/Tools/psa/. Only the C-terminal portion of the alignment is shown, where the key features of interest are found (see Fig. S1 for full alignment). Box 1. Residues involved in binding to the −10 promoter region (region 2). Box 2. Residues involved in binding to the −35 promoter region (region 4.2). Underlined are the two residues in the *E. coli* sigma 70 factor RpoD, K593 and R599, required for interaction with two residues of RhaS (D250 and D241 respectively) and the conserved residues found in the *Synechocystis* ortholog (R412 and K406 respectively).

### L-rhamnose is not metabolised by nor toxic to *Synechocystis*

Before testing whether the *E. coli rhaBAD* promoter is functional in *Synechocystis*, we first wanted to check if the natural sugar inducer L-rhamnose was metabolised by the cyanobacterium or if the use of a non-metabolisable analog of rhamnose would be required, as previously found in *E. coli*^50^. Wild-type *Synechocystis* cells were cultivated in BG11 medium under constant light, with L-rhamnose added to the culture to a final concentration of 1 mg/ml L-rhamnose or omitted in the control. The concentration of L-rhamnose in the culture supernatant was monitored over time using HPLC-RID (Figure 3A). The concentration of L-rhamnose does not change over the course of the experiment, indicating that it is not metabolised by *Synechocystis* in photoautotrophic conditions, nor degraded by exposure to light. Next the effect of L-rhamnose on growth was investigated by monitoring the optical density (OD) at 750 nm of cultures over time, with or without L-rhamnose (Figure 3B). No negative effect of L-rhamnose on growth was observed indicating that L-rhamnose is not inhibitory to *Synechocystis* growth.

**Figure 3.**
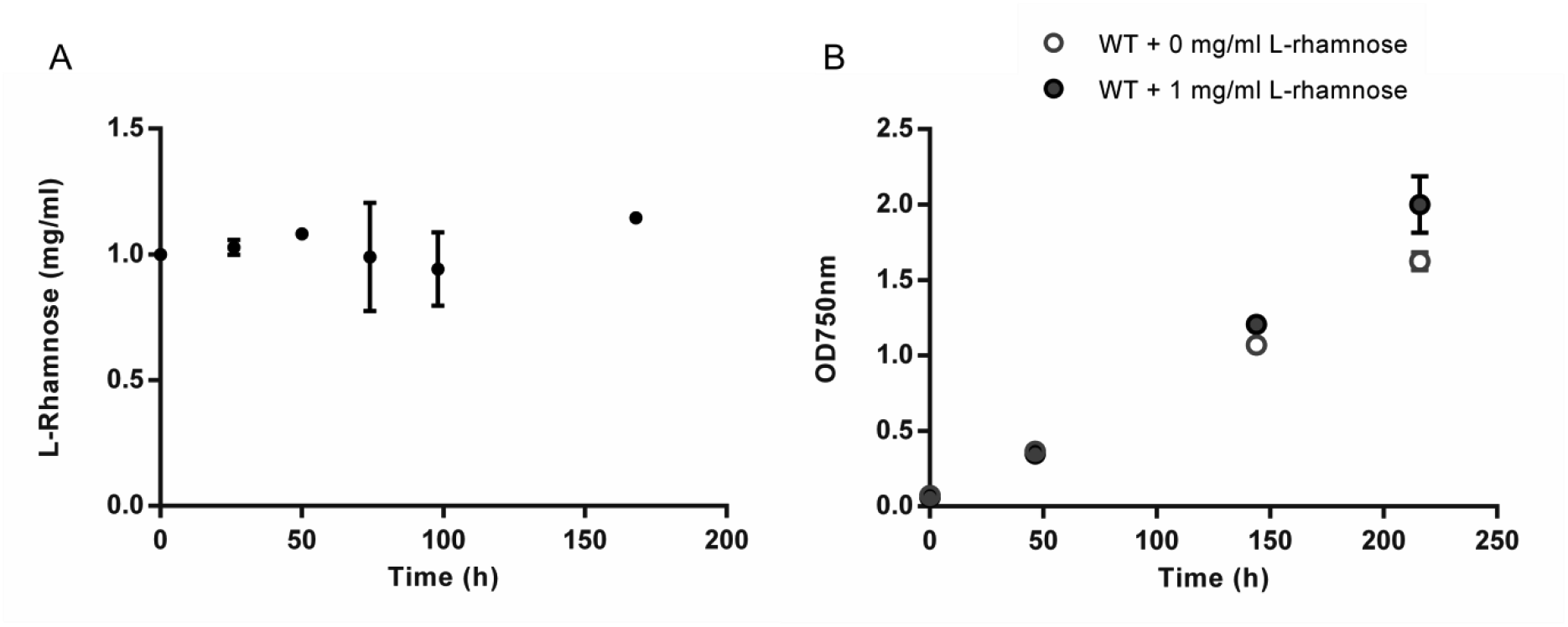
L-rhamnose stability and impact on growth in photoautotrophic cultures of *Synechocystis*. A. Concentration of L-rhamnose over time in the supernatant of photoautotrophic cultures of wild-type *Synechocystis*, as measured by HPLC-RID. B. Growth of wild-type *Synechocystis* in photoautotrophic conditions with and without 1 mg/ml L-rhamnose. Error bars represent the standard deviation of three independent biological replicates.

### Characterisation of *rhaBAD* promoter system in *Synechocystis*

To facilitate the testing of the *rhaBAD* promoter from *E. coli* in *Synechocystis*, an *E. coli-Synechocystis* shuttle reporter plasmid pCK306 (Figure 4) containing the *rhaBAD* promoter sequence and the *rhaS* gene encoding its transcriptional activator was constructed (see Plasmid Construction section of Supplementary Information for details). This plasmid contains homology arms for integration into the genome of *Synechocystis* at the ssl0410 locus, the p15A origin of replication for *E. coli*, the promoter of the *rhaBAD* operon from *E. coli* (P_*rhaBAD*_), a reporter gene encoding yellow fluorescent protein (YFP), a kanamycin-resistance gene functional in both *Synechocystis* and *E. coli*, and *rhaS* from *E. coli*, which encodes the transcriptional activator of the *rhaBAD* promoter, RhaS.. In this genetic context, the native *E. coli* RBS of *rhaS* was predicted to have a T.I.R. of just 72^68^. To determine whether it is necessary to supply *rhaS* heterologously, a control reporter plasmid, pCK305, identical to pCK306 except lacking *rhaS*, was also constructed.

**Figure 4.**
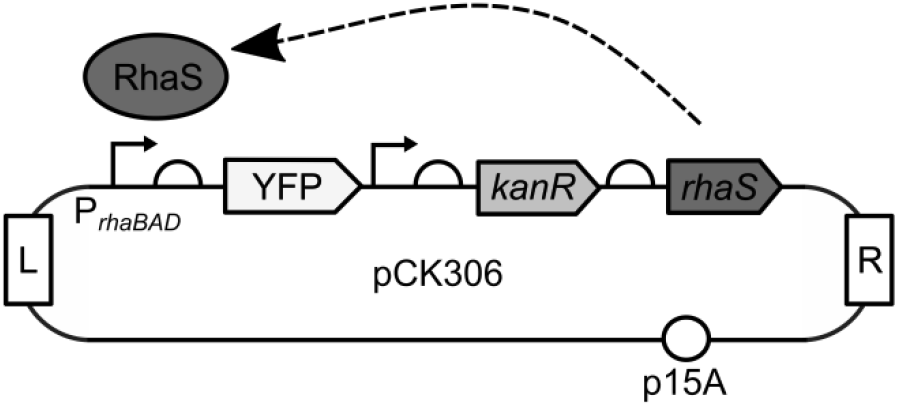
*E. coli-Synechocystis* shuttle plasmid pCK306 containing a YFP reporter of the *rhaBAD* promoter. L and R denote left and right homology arms respectively for integration into the *Synechocystis* sp. PCC 6803 genome at a previously-used insertion site within the ssl0410 ORF adjacent to the *ndhB* locus^69^. *P_rhaBA_D* is the *rhaBAD* promoter sequence from *E. coli*. YFP encodes yellow fluorescent protein. *kanR* encodes an aminoglycoside phosphotransferase which confers resistance to kanamycin in both *Synechocystis* and *E. coli. rhaS* encodes the transcriptional activator of *rhaBAD* promoter, RhaS. p15A is the medium-copy origin of replication which allows replication of the shuttle plasmid in *E. coli*.

To test for L-rhamnose induction of the *rhaBAD* promoter in *Synechocystis*, wild-type cells were transformed with either pCK305 or pCK306 and kanamycin-resistant transformants were passaged until complete segregation was confirmed by PCR. These transformants were then cultured under constant light in BG11 media supplemented with kanamycin, with or without glucose. Cultures were adjusted to a starting optical density (measured at 750 nm) of 0.1, grown for 24 h and then L-rhamnose was added to a range of final concentrations. To determine the response of the promoter to the concentration of the inducer L-rhamnose, the fluorescence intensity of each cell was measured using flow cytometry after 116 h of growth for both photoautotrophic and mixotrophic cultures (Figure 5A & C). Cell density was monitored during growth by measuring optical density of cultures at 750 nm. At the time of sampling, cultures were in the mid-exponential phase of growth. Small differences in optical density were observed between cultures containing glucose and those without glucose. Fluorescence intensity of individual cells (10,000 cells per sample) was measured by flow cytometry, avoiding the need to normalise the fluorescence intensity of culture volumes by optical density, which can be problematic as highly pigmented cyanobacterial cells can partially quench fluorescence. Cells containing the reporter plasmid pCK305, lacking *rhaS*, were unresponsive to any concentration of L-rhamnose added, whereas cells containing the plasmid constitutively expressing *rhaS*, pCK306, show a linear response in YFP fluorescence to increasing concentrations of L-rhamnose in both photoautotrophic and mixotrophic conditions. Saturation of induction occurs at lower concentrations in mixotrophic conditions (0.6 mg/ml) than photoautotrophic conditions (no saturation at 1 mg/ml). To determine the kinetics of YFP expression from the *rhaBAD* promoter in *Synechocystis*, the fluorescence intensity of cells sampled from in the same transformant cultures was monitored over a longer period (Figure 5B & D). Fluorescence was observed in cells containing pCK306 after only 24 h of induction and showed sustained induction in both photoautotrophic and mixotrophic growth conditions, with no decrease in fluorescence observed after > 250 h of growth. Finally, as levels of gene expression can differ among cells in a population of either in natural or engineered strains, flow cytometry was used to investigate the modality (distribution) of fluorescence across *Synechocystis* cells containing pCK306. Induction of the *rhaBAD* promoter in *Synechocystis* containing pCK306 in photoautotrophic conditions was unimodal at all time points measured (Figure S2A). In mixotrophic conditions at the early stages of induction (120 h), some bimodality was observed (Figure S2B), with 3-6% of cells failing to be induced at this time point, however when induction was complete at a later time point (215 h) the induction was unimodal once again (Figure S2C). These data demonstrate that the *rhaBAD* promoter from *E. coli* is functional in *Synechocystis*, allows the strength of expression of a gene of interest to be precisely controlled in both phototrophic and mixotrophic growth conditions and that the transcriptional activator RhaS from *E. coli* is required for function in *Synechocystis*.

**Figure 5.**
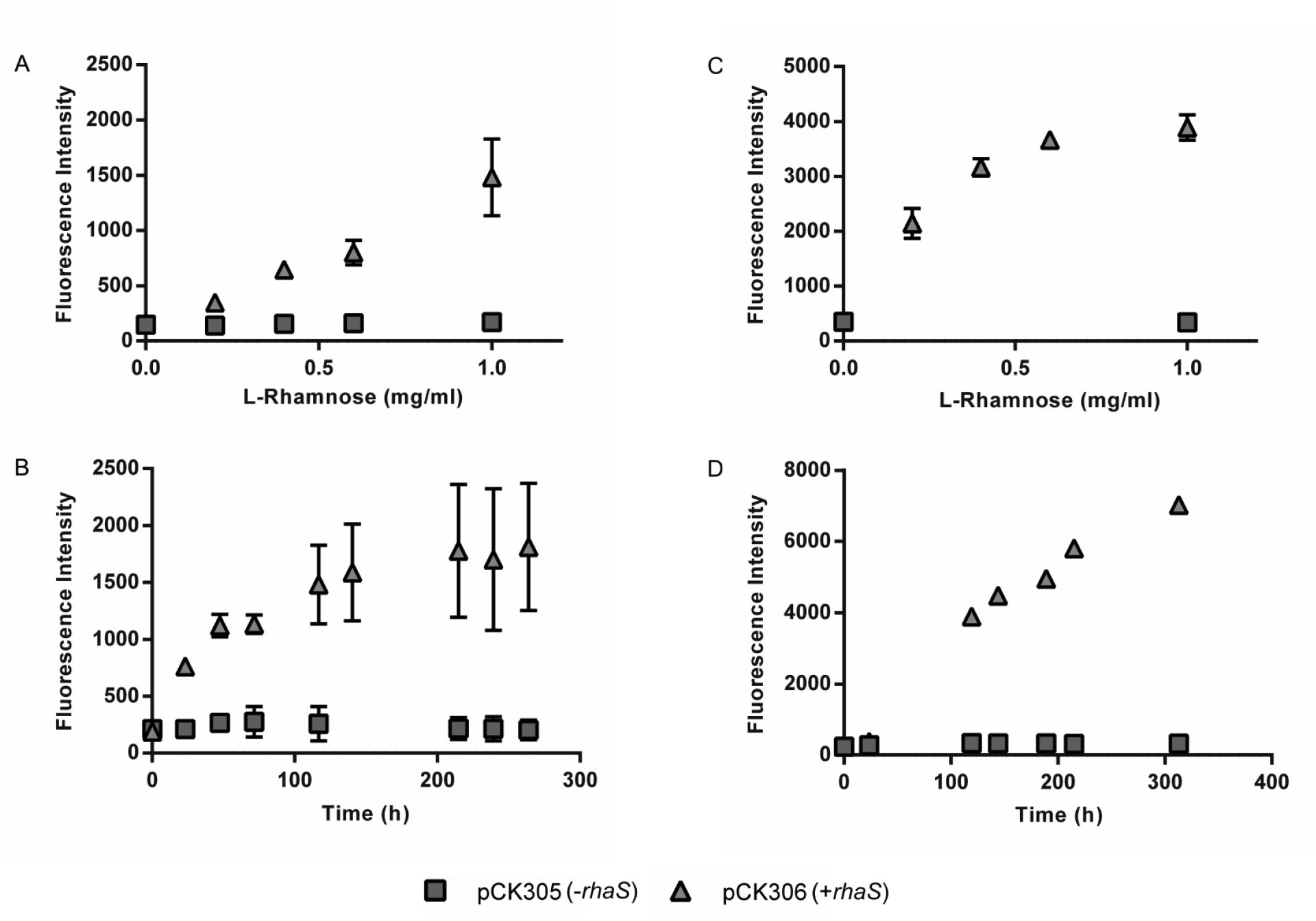
Response to concentration of inducer L-rhamnose and timecourse of induction of *rhaBAD* promoter in *Synechocystis*. A. *Synechocystis* cells containing either pCK305 (*rhaBAD* promoter and YFP only) or pCK306 (*rhaBAD* promoter, YFP and *rhaS*) were cultured in BG11 media supplemented with specified concentrations of L-rhamnose in photoautotrophic conditions and the fluorescence intensity of 10,000 cells measured after 116 h using flow cytometry. B. The same strains of *Synechocystis* were cultured in BG11 media supplemented with L-rhamnose to a final concentration of 1 mg/ml in photoautotrophic conditions and the fluorescence intensity of 10,000 cells measured at specified timepoints using flow cytometry. C. Equivalent experiment to A but strains cultured in BG11 supplemented with 5 mM D-glucose (mixotrophic growth). D. Equivalent experiment to B but strains cultured in BG11 supplemented with 5mM D-glucose (mixotrophic). Error bars shown are the standard deviation of the mean for three independent biological replicates.

Having confirmed that the *rhaBAD* promoter was functional in *Synechocystis* and demonstrated many of the desired properties of an ideal inducible promoter system, we next investigated if modifications to the promoter sequence itself or varying the concentration of RhaS in the cell affected the behaviour of the system. As the role of CRP is still poorly understood in *Synechocystis* and as the CRP-binding site is required for *rhaBAD* functioning in *E. coli*, we investigated the effect that deleting this operator sequence from the promoter would have on induction strength and/or kinetics. The reporter plasmids pCK305 and pCK306 were both modified through deletion of the CRP-binding operator sites, resulting in pCK313 and pCK314 respectively. Wild-type *Synechocystis* cells were transformed with either plasmid and integration and segregation confirmed as before. Transformants were then cultured in both photoautotrophic and mixotrophic growth conditions and the inducer-response and timecourse experiments repeated (Figure 6). Results were very similar to those observed with pCK305 and pCK314, meaning the CRP-binding site is not required for the *rhaBAD* promoter to function in *Synechocystis*.

**Figure 6.**
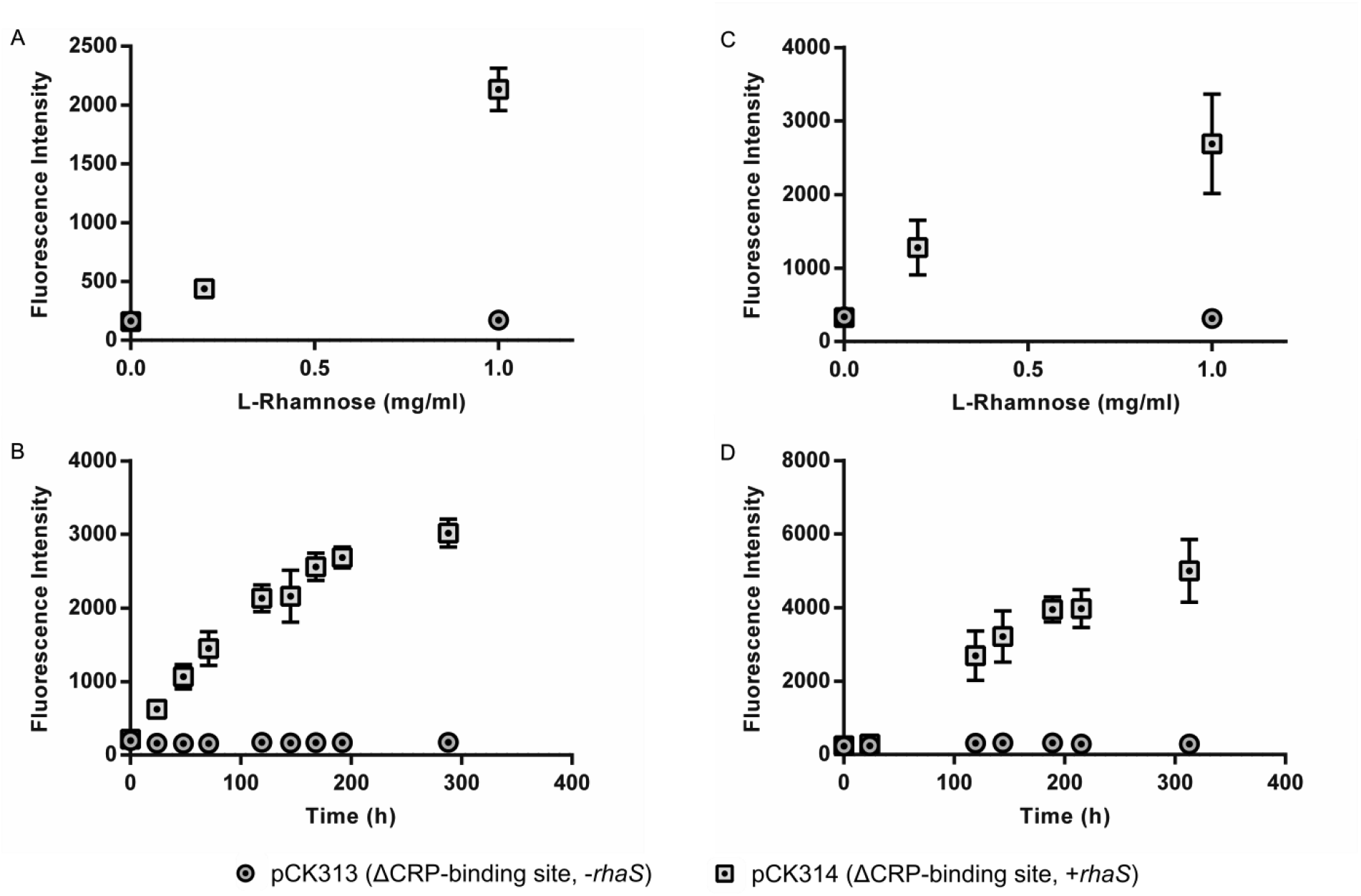
Response to concentration of inducer L-rhamnose and timecourse of induction of a variant of the *rhaBAD* promoter with CRP-binding site deletion. A. *Synechocystis* cells containing either pCK313 (*rhaBAD* promoter minus CRP-binding site and YFP only) or pCK314 (*rhaBAD* promoter minus CRP-binding site, YFP and *rhaS*) were cultured in BG11 media supplemented with specified concentrations of L-rhamnose in photoautotrophic conditions and fluorescence intensity of 10,000 cells measured after 116 h using flow cytometry. B. The same strains of *Synechocystis* were cultured in BG11 media supplemented with L-rhamnose to a final concentration of 1 mg/ml in photoautotrophic conditions and the fluorescence intensity of 10,000 cells measured at specified timepoints using flow cytometry. C. Equivalent experiment to A but strains cultured in BG11 supplemented with 5 mM D-glucose (mixotrophic growth). D. Equivalent experiment to B but strains cultured in BG11 supplemented with 5 mM D-glucose (mixotrophic). Error bars shown are the standard deviation of the mean for three independent biological replicates.

Next we investigated whether increasing the cellular concentration of the transcriptional activator RhaS would change the response to inducer concentration, dynamic range or kinetics of *rhaBAD* promoter induction. The original *rhaS* RBS was predicted to have a low T.I.R. of just 72, so two synthetic RBSs were designed using the RBS Calculator ^68^ with much higher T.I.R. values of 5,000 and 18,000, and these new RBS sequences were inserted in place of the *rhaS* RBS used in pCK306, resulting in pCK320 and pCK321 respectively. These constructs were introduced into *Synechocystis*, integration and complete segregation was confirmed as before, then these transformants were used for inducer-response and timecourse experiments as before. The *Synechocystis* strains transformed with the new RBS variant plasmids pCK320 or pCK321 showed similar fluorescence response to inducer concentration and timecourses to cells transformed with pCK306 (Figure S1).

Finally, we sought to directly compare all the functional *rhaBAD* expression system variants. Absolute levels of fluorescence measured using flow cytometry cannot be directly compared between different days and experiments due to instrument variation. This is sometimes overcome in reporter studies by normalising to a reference promoter included in each separate experiment, allowing relative comparisons. Here, as we had a defined set of constructs to compare, we compared these directly in a single experiment. *Synechocystis* cells containing each of the rhaBAD-promoter reporter plasmids were cultured, both photoautotrophically and mixotrophically, in BG11 media supplemented with 1 mg/ml L-rhamnose, and the fluorescence intensity measured by flow cytometry after 191 h (Figure S4). No statistically-significant difference was observed between cells containing constructs pCK306 (+rhaS), pCK314 (*+rhaS*, ΔCRP-binding site), pCK320 (+rhaS, T.I.R. of RBS of *rhaS* = 5,000) or pCK321 (*+rhaS*, T.I.R. of RBS of *rhaS* = 18,000).

The inducible reporter constructs described above show non-zero levels of fluorescence in *Synechocystis* even in the complete absence of inducer, which could suggest that the promoter is ‘leaky’. However, it was noted that even cells containing the non-functional promoter reporter constructs (such as pCK305) were slightly more fluorescent than wild type cells lacking any reporter plasmid (Figure 7A). As these constructs are integrated into the *Synechocystis* genome, it was hypothesised that this basal fluorescence resulted from transcriptional read through from the chromosome rather than leaky expression from the *rhaBAD* promoter itself. To test this hypothesis, the *rhaBAD* promoter of pCK321 (one of the above-described derivatives of pCK306 which performs identically) was removed resulting in the promoterless plasmid pCK324. This construct was integrated into the same site on the *Synechocystis* genome as all other reporter plasmids, fully segregated and the timecourse experiments in mixotrophic and photoautotrophic growth conditions performed as before. Cells containing pCK324, lacking the *rhaBAD* promoter had the same level of basal YFP fluorescence whether L-rhamnose was added to the media or not and the level of fluorescence in both cases was the same as cells containing pCK305 or pCK306 without inducer. This confirmed that chromosomal read-through was the cause of basal YFP fluorescence and the *rhaBAD* promoter itself was not leaky in the absence of inducer.

**Figure 7.**
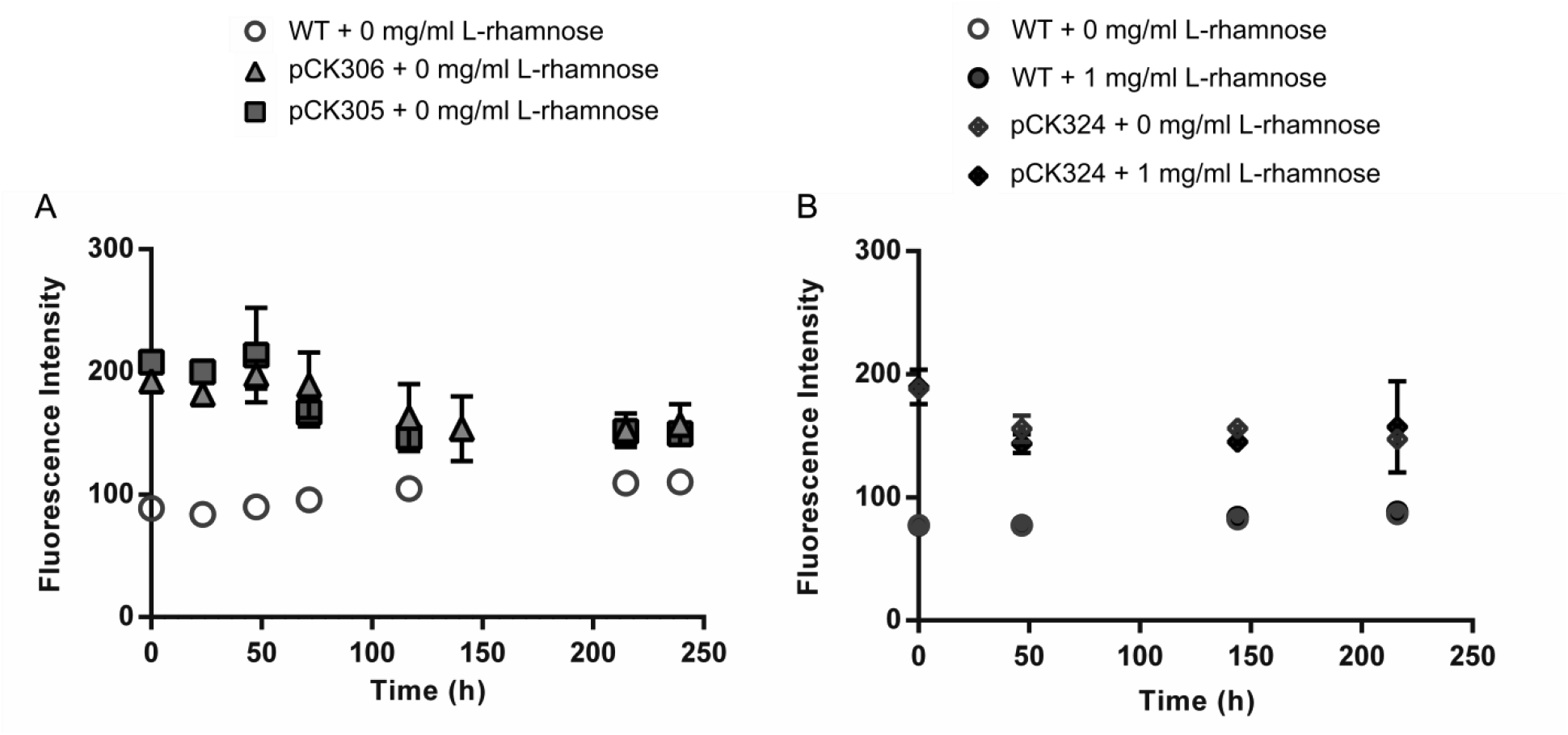
Chromosomal read-through from the site of *Synechocystis* genome integration is responsible for the low basal level of YFP fluorescence observed with *rhaBAD* reporter plasmids. A. Wild-type *Synechocystis* (WT) or *Synechocystis* containing pCK305 (*rhaBAD* promoter and YFP only) or pCK306 (*rhaBAD* promoter, YFP and *rhaS*) were cultured in BG11 media without L-rhamnose and the fluorescence intensity of 10,000 cells measured by flow cytometry. B. Wild-type *Synechocystis* (WT) or *Synechocystis* containing pCK324 (a control vector lacking P*rhaBAD*) were cultured with or without 1 mg/ml L-rhamnose and the fluorescence intensity of 10,000 cells measured by flow cytometry.

### Conclusions

This study tested and showed that the *E. coli rhaBAD* promoter performs excellently as an inducible promoter in the cyanobacterium *Synechocystis* sp. PCC 6803, with a linear response to inducer concentration, good dynamic range, sustained induction in light over long periods and crucially no basal expression in the absence of inducer. For many *Synechocystis* projects and applications, the use of this promoter should allow more precise control of the timing and strength of expression than alternative cyanobacterial inducible promoters. Heterologous expression of *rhaS* was required for promoter function in *Synechocystis*, which is consistent with the apparent absence of an ortholog in the *Synechocystis* genome. This lack of complementation of the *rhaBAD* promoter system by any native *Synechocystis* protein suggests that the heterologously-supplied transcriptional activator RhaS is unlikely to interact with other *Synechocystis* promoters, providing a useful level of independence (or orthogonality). Deletion of the CRP-binding sites from the *rhaBAD* promoter had no effect on promoter function in *Synechocystis* in the experimental conditions tested, including when glucose was added to the growth media. This was unexpected as the function of the *rhaBAD* promoter in *E. coli* requires binding of CRP. For those interested in using the *rhaBAD* promoter in fundamental studies of the circadian clock or photosynthesis, or in applications where cyanobacteria are grown in light and dark cycles, the use of the CRP-binding site deletion variant pCK314 may result in alternative induction responses, as cAMP levels are known to increase in cyanobacteria at night^55^.

The only observed flaw with this implementation of the *rhaBAD* promoter in *Synechocystis* was a low level of basal expression, which we found was independent of the *rhaBAD* promoter. The *sll0410* insertion site adjacent to *ndhB* has been used previously, but seems to result in transcriptional read-through of inserts, presumably from the promoter found inside the *ndhB* ORF^69^. For most inducible expression studies, this observation will be unimportant and expression constructs reported here will be ideal, because in many cases the ability to specify extremely low expression levels is not required. Where extremely low or zero basal and induced expression is required, alternative integration sites or extrachromosomal plasmids may prove more suitable^70^.

We found that the *rhaBAD* promoter of *E. coli* was functional and inducible in *Synechocystis* without any modification of the promoter sequence itself. This was not obvious in advance given reports of difficulties in using *E. coli* promoters in cyanobacteria. In this case our analysis of the relevant transcription factor machinery and interacting residues successfully predicted function of this promoter in *Synechocystis*, so it is interesting to consider whether this promoter might function in other cyanobacteria such as *Synechococcus* sp. PCC 7002 or *Arthrospira* species. For example, one of the sigma 70 factor residues important for interaction with RhaS, K593, is not found in the *Synechococcus* sp. PCC 7002 ortholog but is found in the *Arthrospira plantensis* ortholog. The residue found in the *Synechococcus* ortholog is an arginine, a similar basic amino acid, so may still interact appropriately with RhaS for function.

This study represents an important step towards addressing the shortage of reliable synthetic biology tools for the manipulation of cyanobacteria, both for fundamental and applied studies. The characteristics of the rhamnose-inducible expression system shown in this work will allow greater control of gene expression in cyanobacteria than previously possible. Despite this progress, much work remains in the development and characterisation of other synthetic biology tools to address the unique challenge of engineering these important photoautotrophic organisms and realising their applied potential.

## MATERIALS AND METHODS

### Bacterial strains and Growth Conditions

*E. coli* strain DH5α was used for all plasmid construction and propagation. *Synechocystis* sp. PCC 6803 (the glucose-tolerant derivative of the wild type, obtained from the Nixon lab at Imperial College London) was used for all cyanobacterial experiments. *E. coli* were routinely cultured in LB at 37 °C with shaking at 240 rpm and *Synechocystis* cultured in TES-buffered (pH 8.2) BG11 media^37^ with 5 mM glucose (mixotrophic growth) or without glucose (photoautotrophic growth) at 30 °C with agitation at 150 rpm, supplemented with 30 μg ml^−1^ kanamycin where required. *Synechocystis* were grown in constant white light at 50 μmol m^−2^ s^−1^.

### Plasmid Construction

A table of all plasmids and oligonucleotides (Table S1) is provided in the Supplementary Information. All plasmid construction was carried out using standard molecular cloning methods. Full details are provided in the Supplementary Information.

### Strain Construction

*Wild-type* Synechocystis cells were cultured in BG11 supplemented with 5 mM D-glucose to an optical density (measured at 750 nm) of 0.5 and 4 ml harvested by centrifugation at 3200 g for 15 mins. Pellets were resuspended in 100 μl BG11, 100 ng of plasmid DNA was added and the mixture was incubated at 100 μmol m^−2^ s^−1^ white light for 60 mins. Cells were spotted onto BG11 glucose plates and incubated at 100 μmol m^−2^ s^−1^ white light for 24 h at 30°C. Cells were collected and transferred onto BG11 glucose plates supplemented with 30 μg ml^-1^ kanamycin. When single colonies appeared, transformants were segregated through passaging on selective plates and full segregation was confirmed by PCR.

### Assays

After confirmation by PCR that *Synechocystis* transformants were fully segregated, cells were cultured to mid exponential phase before subculture to a final optical density (measured at 750 nm) of 0.1. Cultures were grown for 24 h and then L-rhamnose added to a variety of final concentrations. The optical density of cultures was monitored at 750 nm and high-resolution fluorescence intensity of each cell was performed using flow cytometry using an Attune NxT Flow Cytometer (ThermoFisher). Cells were gated using forward and side scatter, and GFP fluorescence (excitation and emission wavelengths: 488 and 525 nm [with 20 nm bandwidth] respectively) was measured. Histograms of fluorescence intensity were plotted, and mean statistics extracted.

## ASSOCIATED CONTENT

Supplementary Information available online

## AUTHOR INFORMATION

### Author Contributions

CK and JH designed the study; CK performed experiments; CK, AH and ATM performed plasmid construction; CK and JH prepared the manuscript with input from AH and ATM.

## Notes

The authors declare no competing financial interest.

## ACKNOWLEDGEMENTS

This work was supported by the Biotechnology and Biological Sciences Research Council [BB/M011321/1 to JTH]. The authors thank Pawel Mordaka, George Taylor, Linda Dekker and Lara Sellés Vidal for useful discussions.

## REFERENCES

(1) Nozzi, N. E., and Atsumi, S. (2015) Genome Engineering of the 2,3-Butanediol Biosynthetic Pathway for Tight Regulation in Cyanobacteria. ACS Synth. Biol. 4, 1197–1204.

(2) Oliver, J. W. K., Machado, I. M. P., Yoneda, H., and Atsumi, S. (2013) Cyanobacterial conversion of carbon dioxide to 2,3-butanediol. Proc. Natl. Acad. Sci. U. S. A. 110, 1249–1254.

(3) Angermayr, S. A., van der Woude, A. D., Correddu, D., Vreugdenhil, A., Verrone, V., and Hellingwerf, K. J. (2014) Exploring metabolic engineering design principles for the photosynthetic production of lactic acid by Synechocystis sp. PCC6803. Biotechnol. Biofuels 7, 99.

(4) Varman, A. M., Xiao, Y., Pakrasi, H. B., and Tang, Y. J. (2013) Metabolic engineering of Synechocystis sp. strain PCC 6803 for isobutanol production. Appl. Environ. Microbiol. 79, 908–914.

(5) Englund, E., Andersen-Ranberg, J., Miao, R., Hamberger, B., and Lindberg, P. (2015) Metabolic Engineering of Synechocystis sp. PCC 6803 for Production of the Plant Diterpenoid Manoyl Oxide. ACS Synth. Biol. 4, 1270–1278.

(6) Deng, M. D., and Coleman, J. R. (1999) Ethanol synthesis by genetic engineering in cyanobacteria. Appl. Environ. Microbiol. 65, 523–528.

(7) Namakoshi, K., Nakajima, T., Yoshikawa, K., Toya, Y., and Shimizu, H. (2016) Combinatorial deletions of glgC and phaCE enhance ethanol production in Synechocystis sp. PCC 6803. J. Biotechnol. 239, 13–19.

(8) Choi, Y.-N., and Park, J. M. (2016) Enhancing biomass and ethanol production by increasing NADPH production in Synechocystis sp. PCC 6803. Bioresour. Technol. 213, 54–57.

(9) Dexter, J., and Fu, P. (2009) Metabolic engineering of cyanobacteria for ethanol production. Energy Environ. Sci. 2, 857–864.

(10) Ranade, S., Zhang, Y., Kaplan, M., Majeed, W., and He, Q. (2015) Metabolic Engineering and Comparative Performance Studies of Synechocystis sp. PCC 6803 Strains for Effective Utilization of Xylose. Front. Microbiol. 6, 1484.

(11) Redinbo, M. R., Yeates, T. O., and Merchant, S. (1994) Plastocyanin: structural and functional analysis. J. Bioenerg. Biomembr. 26, 49–66.

(12) Grotjohann, I., and Fromme, P. (2005) Structure of cyanobacterial photosystem I. Photosynth. Res. 85, 51–72.

(13) Nickelsen, J., and Rengstl, B. (2013) Photosystem II assembly: from cyanobacteria to plants. Annu. Rev. Plant Biol. 64, 609–635.

(14) Mulkidjanian, A. Y., Koonin, E. V., Makarova, K. S., Mekhedov, S. L., Sorokin, A., Wolf, Y. I., Dufresne, A., Partensky, F., Burd, H., Kaznadzey, D., Haselkorn, R., and Galperin, M. Y. (2006) The cyanobacterial genome core and the origin of photosynthesis. Proc. Natl. Acad. Sci. U. S. A. 103, 13126–13131.

(15) Golbeck, J. H. (1994) Photosystem I in Cyanobacteria, in The Molecular Biology of Cyanobacteria, pp 319–360. Springer, Dordrecht.

(16) Heinz, S., Liauw, P., Nickelsen, J., and Nowaczyk, M. (2016) Analysis of photosystem II biogenesis in cyanobacteria. Biochim. Biophys. Acta 1857, 274–287.

(17) Cohen, S. E., and Golden, S. S. (2015) Circadian Rhythms in Cyanobacteria. Microbiol. Mol. Biol. Rev. 79, 373–385.

(18) Golden, S. S., Ishiura, M., Johnson, C. H., and Kondo, T. (1997) CYANOBACTERIAL CIRCADIAN RHYTHMS. Annu. Rev. Plant Physiol. Plant Mol. Biol. 48, 327–354.

(19) Kondo, T., and Ishiura, M. (2000) The circadian clock of cyanobacteria. Bioessays 22, 10–15.

(20) Yeates, T. O., Kerfeld, C. A., Heinhorst, S., Cannon, G. C., and Shively, J. M. (2008) Protein-based organelles in bacteria: carboxysomes and related microcompartments. Nat. Rev. Microbiol. 6, 681–691.

(21) Rae, B. D., Long, B. M., Badger, M. R., and Price, G. D. (2013) Functions, compositions, and evolution of the two types of carboxysomes: polyhedral microcompartments that facilitate CO2 fixation in cyanobacteria and some proteobacteria. Microbiol. Mol. Biol. Rev. 77, 357–379.

(22) Shively, J. M., and English, R. S. (1991) The carboxysome, a prokaryotic organelle: a minireview. Can. J. Bot.

(23) Tanaka, S., Kerfeld, C. A., Sawaya, M. R., Cai, F., Heinhorst, S., Cannon, G. C., and Yeates, T. O. (2008) Atomic-level models of the bacterial carboxysome shell. Science 319, 1083–1086.

(24) Griese, M., Lange, C., and Soppa, J. (2011) Ploidy in cyanobacteria. FEMS Microbiol. Lett. 323, 124–131.

(25) Zerulla, K., Ludt, K., and Soppa, J. (2016) The ploidy level of Synechocystis sp. PCC 6803 is highly variable and is influenced by growth phase and by chemical and physical external parameters. Microbiology 162, 730–739.

(26) Jones, P. R. (2014) Genetic instability in cyanobacteria - an elephant in the room? Front Bioeng Biotechnol 2, 12.

(27) Markley, A. L., Begemann, M. B., Clarke, R. E., Gordon, G. C., and Pfleger, B. F. (2015) Synthetic biology toolbox for controlling gene expression in the cyanobacterium Synechococcus sp. strain PCC 7002. ACS Synth. Biol. 4, 595–603.

(28) Boyanapalli, R., Bullerjahn, G. S., Pohl, C., Croot, P. L., Boyd, P. W., and McKay, R. M. L. (2007) Luminescent whole-cell cyanobacterial bioreporter for measuring Fe availability in diverse marine environments. Appl. Environ. Microbiol. 73, 1019–1024.

(29) Zhou, J., Zhang, H., Meng, H., Zhu, Y., Bao, G., Zhang, Y., Li, Y., and Ma, Y. (2014) Discovery of a super-strong promoter enables efficient production of heterologous proteins in cyanobacteria. Sci. Rep. 4, 4500.

(30) Wilms, B., Hauck, A., Reuss, M., Syldatk, C., Mattes, R., Siemann, M., and Altenbuchner, J. (2001) High-cell-density fermentation for production of L-N-carbamoylase using an expression system based on the Escherichia coli rhaBAD promoter. Biotechnol. Bioeng. 73, 95–103.

(31) Saïda, F., Uzan, M., Odaert, B., and Bontems, F. (2006) Expression of highly toxic genes in E. coli: special strategies and genetic tools. Curr. Protein Pept. Sci. 7, 47–56.

(32) Terpe, K. (2006) Overview of bacterial expression systems for heterologous protein production: from molecular and biochemical fundamentals to commercial systems. Appl. Microbiol. Biotechnol. 72, 211–222.

(33) Englund, E., Liang, F., and Lindberg, P. (2016) Evaluation of promoters and ribosome binding sites for biotechnological applications in the unicellular cyanobacterium Synechocystis sp. PCC 6803. Sci. Rep. 6, 36640.

(34) Blasi, B., Peca, L., Vass, I., and Kós, P. B. (2012) Characterization of stress responses of heavy metal and metalloid inducible promoters in synechocystis PCC6803. J. Microbiol. Biotechnol. 22, 166–169.

(35) Čelešnik, H., Tanšek, A., Tahirović, A., Vižintin, A., Mustar, J., Vidmar, V., and Dolinar, M. (2016) Biosafety of biotechnologically important microalgae: intrinsic suicide switch implementation in cyanobacterium Synechocystis sp. PCC 6803. Biol. Open 5, 519–528.

(36) Peca, L., Kós, P. B., Máté, Z., Farsang, A., and Vass, I. (2008) Construction of bioluminescent cyanobacterial reporter strains for detection of nickel, cobalt and zinc. FEMS Microbiol. Lett. 289, 258–264.

(37) Rippka, R. (1988) Isolation and purification of cyanobacteria. Methods Enzymol. 167, 3–27.

(38) Ludwig, M., Chua, T. T., Chew, C. Y., and Bryant, D. A. (2015) Fur-type transcriptional repressors and metal homeostasis in the cyanobacterium Synechococcus sp. PCC 7002. Front. Microbiol. 6, 1217.

(39) García-Domínguez, M., Lopez-Maury, L., Florencio, F. J., and Reyes, J. C. (2000) A gene cluster involved in metal homeostasis in the cyanobacterium Synechocystis sp. strain PCC 6803. J. Bacteriol. 182, 1507–1514.

(40) Antonkine, M. L., Maes, E. M., Czernuszewicz, R. S., Breitenstein, C., Bill, E., Falzone, C. J., Balasubramanian, R., Lubner, C., Bryant, D. A., and Golbeck, J. H. (2007) Chemical rescue of a site-modified ligand to a [4Fe-4S] cluster in PsaC, a bacterial-like dicluster ferredoxin bound to Photosystem I. Biochim. Biophys. Acta 1767, 712–724.

(41) Huang, H.-H., and Lindblad, P. (2013) Wide-dynamic-range promoters engineered for cyanobacteria. J. Biol. Eng. 7, 10.

(42) Zess, E. K., Begemann, M. B., and Pfleger, B. F. (2016) Construction of new synthetic biology tools for the control of gene expression in the cyanobacterium Synechococcus sp. strain PCC 7002. Biotechnol. Bioeng. 113, 424–432.

(43) Nozzi, N. E., Case, A. E., Carroll, A. L., and Atsumi, S. (2017) Systematic Approaches to Efficiently Produce 2,3-Butanediol in a Marine Cyanobacterium. ACS Synth. Biol.

(44) Niederholtmeyer, H., Wolfstädter, B. T., Savage, D. F., Silver, P. A., and Way, J. C. (2010) Engineering cyanobacteria to synthesize and export hydrophilic products. Appl. Environ. Microbiol. 76, 3462–3466.

(45) Camsund, D., Heidorn, T., and Lindblad, P. (2014) Design and analysis of LacI-repressed promoters and DNA-looping in a cyanobacterium. J. Biol. Eng. 8, 4.

(46) Abe, K., Miyake, K., Nakamura, M., Kojima, K., Ferri, S., Ikebukuro, K., and Sode, K. (2014) Engineering of a green-light inducible gene expression system in Synechocystis sp. PCC6803. Microb. Biotechnol. 7, 177–183.

(47) Egan, S. M., and Schleif, R. F. A Regulatory Cascade in the Induction of rhaBAD.

(48) Giacalone, M. J., Gentile, A. M., Lovitt, B. T., Berkley, N. L., Gunderson, C. W., and Surber, M. W. (2006) Toxic protein expression in Escherichia coli using a rhamnose-based tightly regulated and tunable promoter system. Biotechniques 40, 355–364.

(49) Haldimann, A., Daniels, L. L., and Wanner, B. L. (1998) Use of new methods for construction of tightly regulated arabinose and rhamnose promoter fusions in studies of the Escherichia coli phosphate regulon. J. Bacteriol. 180, 1277–1286.

(50) Kelly, C. L., Liu, Z., Yoshihara, A., Jenkinson, S. F., Wormald, M. R., Otero, J., Estévez, A., Kato, A., Marqvorsen, M. H. S., Fleet, G. W. J., Estévez, R. J., Izumori, K., and Heap, J. T. (2016) Synthetic Chemical Inducers and Genetic Decoupling Enable Orthogonal Control of the rhaBAD Promoter. ACS Synth. Biol. 5, 1136–1145.

(51) Egan, S. M., and Schleif, R. F. (1994) DNA-dependent renaturation of an insoluble DNA binding protein. Identification of the RhaS binding site at rhaBAD. J. Mol. Biol. 243, 821–829.

(52) Yoshimura, H., Hisabori, T., Yanagisawa, S., and Ohmori, M. (2000) Identification and characterization of a novel cAMP receptor protein in the cyanobacterium Synechocystis sp. PCC 6803. J. Biol. Chem. 275, 6241–6245.

(53) Yoshimura, H., Yanagisawa, S., Kanehisa, M., and Ohmori, M. (2002) Screening for the target gene of cyanobacterial cAMP receptor protein SYCRP1. Mol. Microbiol. 43, 843–853.

(54) Wickstrum, J. R., Skredenske, J. M., Kolin, A., Jin, D. J., Fang, J., and Egan, S. M. (2007) Transcription activation by the DNA-binding domain of the AraC family protein RhaS in the absence of its effector-binding domain. J. Bacteriol. 189, 4984–4993.

(55) Hedger, J., Holmquist, P. C., Leigh, K. A., Saraff, K., Pomykal, C., and Summers, M. L. (2009) Illumination stimulates cAMP receptor protein-dependent transcriptional activation from regulatory regions containing class I and class II promoter elements in Synechocystis sp. PCC 6803. Microbiology 155, 2994–3004.

(56) Forcada-Nadal, A., Forchhammer, K., and Rubio, V. (2014) SPR analysis of promoter binding of Synechocystis PCC6803 transcription factors NtcA and CRP suggests cross-talk and sheds light on regulation by effector molecules. FEBS Lett. 588, 2270–2276.

(57) Berg, O. G., and von Hippel, P. H. (1988) Selection of DNA binding sites by regulatory proteins. II. The binding specificity of cyclic AMP receptor protein to recognition sites. J. Mol. Biol. 200, 709–723.

(58) Altschul, S. F., Gish, W., Miller, W., Myers, E. W., and Lipman, D. J. (1990) Basic local alignment search tool. J. Mol. Biol. 215, 403–410.

(59) Camsund, D., and Lindblad, P. (2014) Engineered transcriptional systems for cyanobacterial biotechnology. Front Bioeng Biotechnol 2, 40.

(60) Hakimi, M.-A., Privat, I., Valay, J.-G., and Lerbs-Mache, S. (2000) Evolutionary Conservation of C-terminal Domains of Primary Sigma70-type Transcription Factors between Plants and Bacteria. J. Biol. Chem. 275, 9215–9221.

(61) Paget, M. S. B., and Helmann, J. D. (2003) The sigma70 family of sigma factors. Genome Biol. 4, 203.

(62) Paget, M. S. (2015) Bacterial Sigma Factors and Anti-Sigma Factors: Structure, Function and Distribution. Biomolecules 5, 1245–1265.

(63) Dombroski, A. J., Walter, W. A., Record, M. T., Jr, Siegele, D. A., and Gross, C. A. (1992) Polypeptides containing highly conserved regions of transcription initiation factor sigma 70 exhibit specificity of binding to promoter DNA. Cell 70, 501–512.

(64) Dombroski, A. J., Walter, W. A., and Gross, C. A. (1993) Amino-terminal amino acids modulate sigma-factor DNA-binding activity. Genes Dev. 7, 2446–2455.

(65) Lonetto, M., Gribskov, M., and Gross, C. A. (1992) The sigma 70 family: sequence conservation and evolutionary relationships. J. Bacteriol. 174, 3843–3849.

(66) Wickstrum, J. R., and Egan, S. M. (2004) Amino Acid Contacts between Sigma 70 Domain 4 and the Transcription Activators RhaS and RhaR. J. Bacteriol. 186, 6277–6285.

(67) Rice, P., Longden, I., and Bleasby, A. (2000) EMBOSS: the European Molecular Biology Open Software Suite. Trends Genet. 16, 276–277.

(68) Salis, H. M., Mirsky, E. A., and Voigt, C. A. (2009) Automated design of synthetic ribosome binding sites to control protein expression. Nat. Biotechnol. 27, 946–950.

(69) Ogawa, T. (1991) A gene homologous to the subunit-2 gene of NADH dehydrogenase is essential to inorganic carbon transport of Synechocystis PCC6803. Proc. Natl. Acad. Sci. U. S. A. 88, 4275–4279.

(70) Pinto, F., Pacheco, C. C., Oliveira, P., Montagud, A., Landels, A., Couto, N., Wright, P. C., Urchueguía, J. F., and Tamagnini, P. (2015) Improving a Synechocystis-based photoautotrophic chassis through systematic genome mapping and validation of neutral sites. DNA Res. 22, 425–437.

